# Proximity interaction analysis of the *Plasmodium falciparum* putative ubiquitin ligase *Pf*RNF1 reveals a role in RNA regulation

**DOI:** 10.1101/2023.03.16.533063

**Authors:** Afia Farrukh, Jean Pierre Musabyimana, Ute Distler, Stefan Tenzer, Gabriele Pradel, Che Julius Ngwa

## Abstract

Some proteins have acquired both ubiquitin ligase activity and RNA-binding properties and are therefore known as RNA-binding Ubiquitin ligases (RBULs). These proteins provide a link between the RNA metabolism and the ubiquitin proteasome system (UPS). The UPS is a crucial protein surveillance system of eukaryotes primarily involved in the selective proteolysis of proteins which are covalently marked with ubiquitin through a series of steps involving ubiquitin E1 activating, E2 conjugating and E3 ligating enzymes. The UPS also regulates other key cellular processes such as cell cycle, proliferation, cell differentiation, transcription and signal transduction. While RBULs have been characterized in other organisms, little is known about their role in *Plasmodium falciparum*, the causative agent of the deadliest human malaria, malaria tropica. In this study, we characterized a previously identified putative *P. falciparum* RING finger E3 ligase *Pf*RNF1. We show that the protein is highly expressed in sexual stage parasites and mainly present in immature male gametocytes. Using proximity interaction studies with parasite lines expressing *Pf*RNF1 tagged with the Biotin ligase BirA, we identified an interaction network of *Pf*RNF1 in both the asexual blood stages and gametocytes composed mainly of ribosomal proteins, RNA-binding proteins including translational repressors such DOZI, CITH, PUF1 and members of the CCR4-NOT complex, as well as proteins of the UPS such as RPN11, RPT1 and RPT6. Our interaction network analysis reveals *Pf*RNF1 as a potential RNA-binding E3 ligase which links RNA dependent processes with protein ubiquitination to regulate gene expression.

**Importance:** RBULs provide a link between RNA-mediated processes with the ubiquitin system. Only a few RBULs have been identified and none has been characterized in the malaria parasite *P. falciparum*. In this study, we unveiled the interactome of the putative *P. falciparum* E3 ligase *Pf*RNF1. We show that *Pf*RNF1 interacts with both proteins of the ubiquitin system as well as RNA-binding proteins therefore indicating that it is a putative RBUL which links RNA regulation with the ubiquitin system in *P. falciparum*.

## INTRODUCTION

Malaria remains one of the most devastating parasitic diseases worldwide with approximately 247 million infections and 619,000 deaths in 2021(1).The disease is caused by protozoa parasites of the genus *Plasmodium* with *Plasmodium falciparum* resulting in malaria tropica being the deadliest form of the human parasite. Its complex life cycle requires two host, the human and the anopheline mosquito where the parasite undergoes several morphological and development stages. Successful development of the parasite through these different morphological and development forms in both hosts requires a very tight regulation of cellular processes.

Post-transcriptional regulation of RNA is one of the most crucial mechanisms of RNA homeostasis and gene regulation. In the malaria parasite, several post-transcription mechanisms of gene regulation have been reported. Proteins associated with the deadenylation-mediated mRNA decay pathway, a major pathway resulting in RNA deadenylation followed by decapping and degradation have been detected in *Plasmodium* including mRNA-decapping enzymes and members of the CCR4-Not complex (2–5). The CCR4-NOT complex is a multi-subunit complex present in all eukaryotes that contributes to regulate gene expression of mRNA (6). A *P. falciparum* deadenylase enzyme caf1 is critical in the regulation of genes associated with parasite egress and invasion (7). In *P. yoelii*, two members of the CAF1/CCR4/NOT complex *Py*NOT1-G and *Py*CCR4-1 have been shown to regulate mRNA important for gametocyte development and transmission (8, 9).

Another important mechanism of post-transcriptional regulation reported in the malaria parasite is translational repression. In this process, transcripts are stored in a translationally repressed state by binding to RNA-binding proteins that condense the transcripts as messenger ribonucleoprotein (mRNP) complexes in cytoplasmic granules. The mRNA is only released when it is needed for parasite development. Translational repression was first demonstrated in the rodent malaria model *P. berghei* where it was shown to store female gametocyte transcripts important for zygote-to-ookinete development such as *Pb*25 and *Pb2*8 in a ribonucleoprotein complex composed of RNA-binding proteins like DOZI (development of zygote inhibitor) and CITH (CAR-I and fly Trailer Hitch). The repressed transcripts are only released after gametocyte activation to allow for their translation to yield proteins important for development of the parasite in the mosquito vector (10, 11). Subsequent studies in *P. falciparum* confirmed the presence of translational repression with the involvement of the Pumilio/Fem-3 binding factor (Puf) family Puf2 and other interaction partners like 7-helix-1 in the translational repression of female gametocyte-specific genes and their storage in stress granules allowing for their release after gametocyte activation (12, 13).

The functional inhibition of some RNA-binding proteins correlates with cancer, autoimmune and neurological diseases (14) and some possess both RNA binding properties and ubiquitin ligase activity and are known as RNA-binding E3 ubiquitin ligases (RBULs). These RBULs therefore link RNA-dependent processes with the ubiquitin proteasome system (UPS) to regulate gene expression (15). The UPS is a major intracellular protein degradation system initiated by a signal cascade whereby ubiquitin is activated by an ubiquitin-activating enzyme E1 in a reaction that requires ATP. The activated ubiquitin is then transferred to ubiquitin conjugating enzyme (E2) and it is finally conjugated to lysine side chains of substrate proteins with the help of the ubiquitin ligase E3 (16, 17). The polyubiquitinated proteins are subsequently transported to the proteasome for degradation. Apart from protein degradation, the addition of ubiquitin or ubiquitin-like proteins to substrates may lead to the regulation of other cellular processes including DNA repair, transcription, cell division, endocytosis and immune response (18–20). RBULs usually have domains which bind RNA such as RRM, CCCH, KH, and Lys-rich domains or E3 ligase domains such as the RING (Really Interesting New Gene) domain. However, some RBULs have been reported to lack a well-defined RNA binding domain but possess only a RING-finger domain, like ARIH2 (21). RBULs have been characterized in other organisms, for example, human MEX-3C is an RBUL, which regulates the expression levels of the gene encoding HLA-A2,a major histocompatibility complex I receptor by binding to the 3’UTR of HLA-A2 mRNA using its KH domain to induce its RING dependent degradation (22). NOT4 is a member of the deadenylation machinery and is also an RBUL implicated in the regulation of transcription in yeast (23).

Previous transcriptional profiling in *P. falciparum* identified a RING-finger domain protein termed *Pf*RNF1, which is epigenetically regulated during gametocyte development (24). The role of *Pf*RNF1 in gametocytes, however, has not been unveiled so far. In this study, we used immunochemical characterization coupled with BioID-based interaction studies to show that that *Pf*RNF1 is a potential RBUL that links ubiquitin-dependent pathways and RNA-binding to regulate gene expression during gametocyte development and transmission.

## RESULTS

### *Pf*RNF1 shows peak transcript and protein expression in immature gametocytes

*Pf*RNF1 is a 135-kDa protein with a RING zinc finger domain (Fig. 1A). We first performed a semi-quantitative RT-PCR using RNA from different asexual blood stage parasites (asexual blood stages (ABSs); rings, trophozoites and schizonts) and immature and mature gametocytes as well as gametocytes at 30 min post-activation to determine the transcript expression. To this end, an equal amount of purified RNA was reverse-transcribed to cDNA and *pfrnf1* transcript was amplified using specific primers (Fig. 1B). The stage-specificity of the samples was verified by using primers specific for either *pfama1*, encoding a protein primarily present in merozoites (25), and for *pfccp2*, encoding the gametocyte-specific LCCL-domain protein *Pf*CCP2 (26). Transcript analysis of the gene *pffbpa* encoding fructose bisphosphate aldolase using specific primers served as positive control (27). Potential gDNA contamination was excluded by using RNA samples lacking reverse transcriptase in combination with *pffbpa*-specific primers. The semi-quantitative RT-PCR analyses showed highest transcript expression of *pfrnf1* in immature gametocytes with very low transcript expression detected in the ABSs (Fig. 1B).

**Figure 1:**
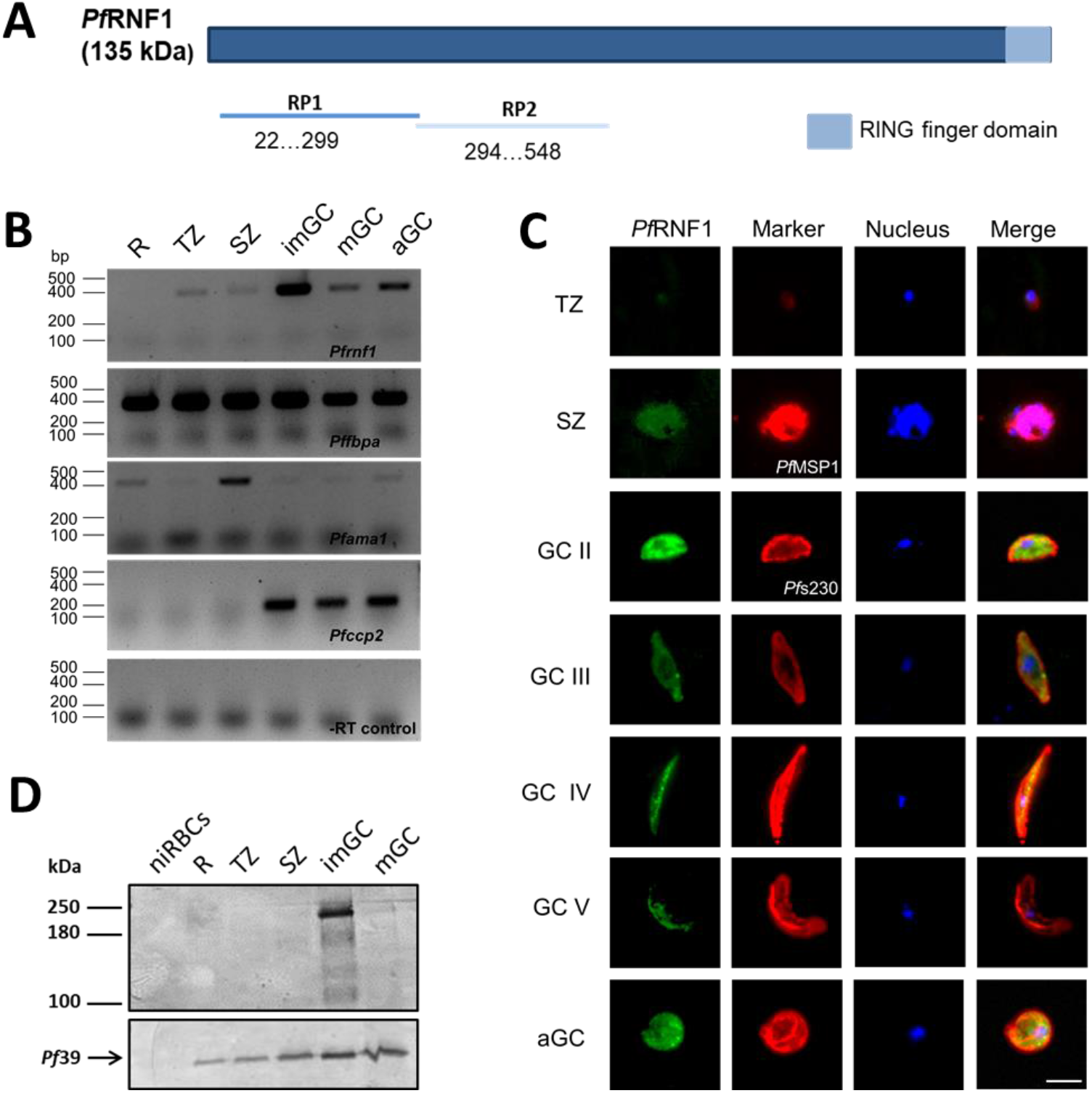
Expression and localization of *Pf*RNF1 in the *P. falciparum* blood stages. (A) Schematic depicting the *Pf*RNF1 domain structure. RP1 and RP2 represent the regions of the recombinant peptide. (B) Transcript expression of *Pf*RNF1 in the blood stages of *P. falciparum*. Diagnostic RT-PCR was performed on mRNA isolated from rings (R), trophozoites (TZ), schizonts (SZ), immature (imGC), mature (mGC) and gametocytes at 30 min post-activation (aGC) to determine the expression levels of *pfrnf1* (398 bp). The expression of *pfama1* (407 bp) and *pfccp2* (198 bp) were used to validate the stage specificity of the samples. Samples lacking reverse transcriptase (−RT) were used to investigate potential contamination with gDNA. Transcript analysis of *pffbpa* (378 bp) was used as positive control. (C) Immunolocalization of *Pf*RNF1 in the blood stages of *P. falciparum* using antisera against RP2. Mouse anti-*Pf*RNF1 was used to immunolabel TZ, SZ and gametocytes (GC) of stages II-V and aGC (green). TZ and SZ were counterlabelled with rabbit anti-*Pf*MSP1 antibody and GCs with rabbit anti-*Pf*s230 antisera (red); nuclei were highlighted by Hoechst 33342 nuclear stain (blue). Bar, 5 μm. (D) *Pf*RNF1-HA expression in blood stage parasites. Lysates from R, TZ, SZ, imGC and mGC of the *Pf*RNF1-HA parasite line were immunoblotted using rabbit anti-HA antibody to detect *Pf*RNF1. Lysate of non-infected RBCs (niRBC) was used as negative control. Equal loading was confirmed using mouse anti-*Pf*39 antisera (∼39 kDa).

The expression of *Pf*RNF1 was then analysed at the protein level. In addition to an antibody previously generated against the recombinant peptide RP1 (24), an antibody against a second peptide, RP2, was produced in mice (Fig. 1A). Both antisera were used to immunolabel *Pf*RNF1 in ABSs and gametocytes. Indirect immunofluorescence assays (IFAs) demonstrated high *Pf*RNF1 levels in the cytoplasm and nucleus of gametocytes with peak expression in immature stage II gametocytes (Fig. 1C). Further, low levels of *Pf*RNF1 were detectable in schizonts. The gender-specific expression of *Pf*RNF1 was investigated by co-immunolabelling of gametocytes with antisera against *Pf*RNF1 in combination with either anti-*Pf*s230 antibody, which labels all gametocytes, or with anti-*Pf*s25 antibody to highlight female gametocytes. A total of 100 gametocytes positive for either-*Pf*s230 or *Pf*s25 were evaluated for a *Pf*RNF1 signal. Quantification revealed that the majority of *Pf*RNF1-positive cells were negative for *Pf*s25, indicating that *Pf*RNF1 is mainly expressed in male gametocytes (Fig. S1). Also, using Western blotting, a high protein expression in immature gametocytes was confirmed using a previously generated *Pf*RNF1-HA (Fig. 1D). *Pf*RNF1 was running at a higher molecular weight of approximately 200 kDa, as has been described before (24).

### *Pf*RNF1-GFP-BirA expression results in protein biotinylation in ABSs and gametocytes

To identify the *Pf*RNF1 interactome, we generated lines episomally expressing *Pf*RNF1-GFP-BirA fusion protein under the control of the ABS promoter *pfama1* and a gametocyte-specific promoter *pffnpa* by transfecting parasites with constructed pARL-*Pf*RNF1-*pfama1*-GFP-BirA and pARL-*Pf*RNF1-*pffnpa*-GFP-BirA vectors respectively (Fig. S2A, (28)). The presence of the episomal vector in the respective transfected lines was confirmed by diagnostic PCR (Fig. S2B). Western blot analysis using anti-GFP antibody confirmed the expression of the *Pf*RNF1-GFP-BirA fusion protein in ABS and gametocyte lysates in the *Pf*RNF1-*pfama1*-GFP-BirA and the *Pf*RNF1-*pffnpa*-GFP-BirA lines, respectively. A protein band of approximately 280 kDa was detected in gametocyte lysate of line *Pf*RNF1-*pffnpa*-GFP-BirA and ABS lysate of the *Pf*RNF1-*pfama1*-GFP-BirA line (Fig. 2A). No protein bands were detected in the wildtype strain NF54 (WT NF54) control (Fig. 2A). IFA using anti-GFP antibody, confirmed of the presence of GFP-tagged *Pf*RNF1 in schizonts and gametocytes, respectively (Fig. 2B).

**Figure 2:**
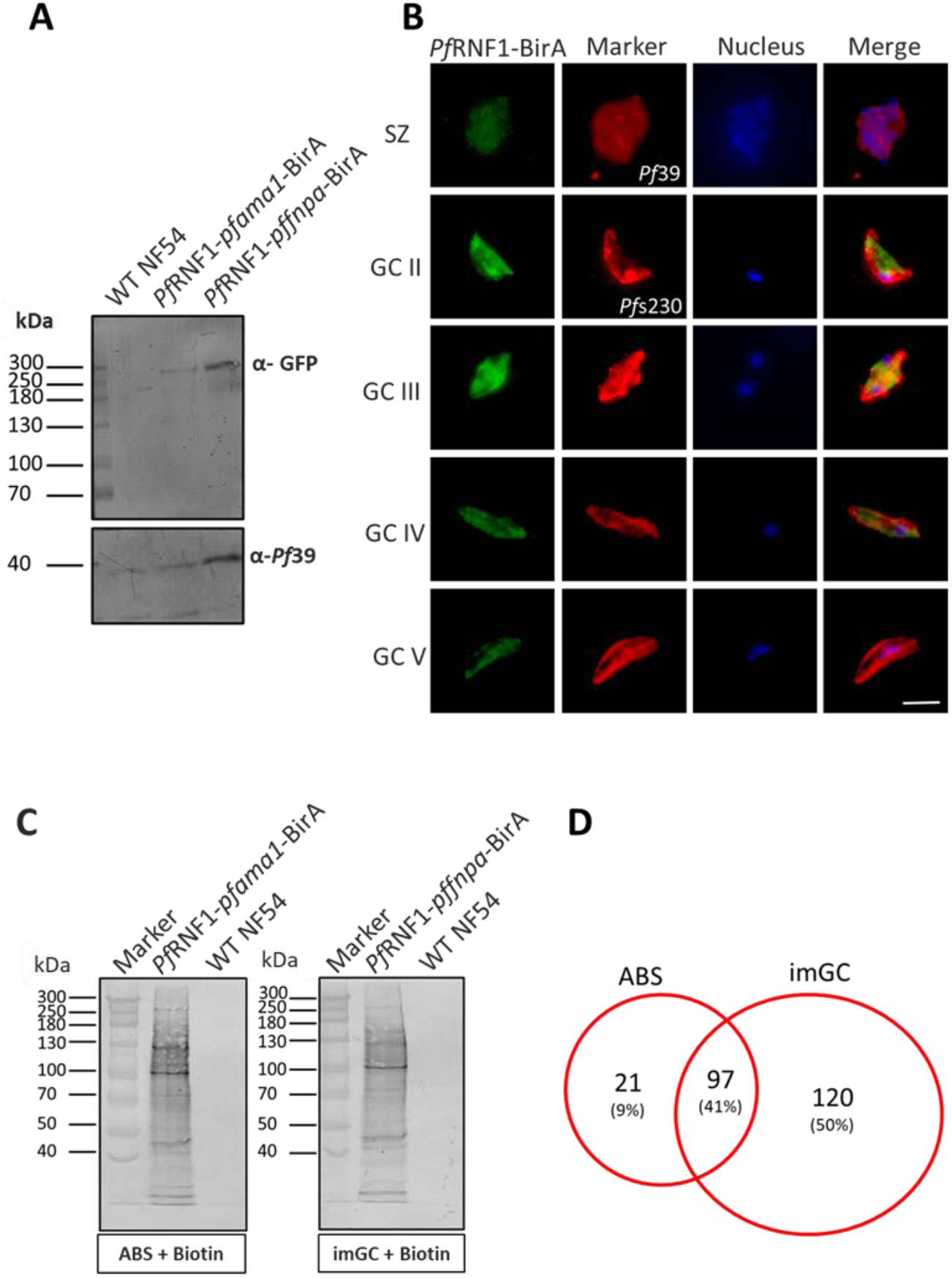
Protein biotinylation in the blood stages of the *Pf*RNF1-GFP-BirA transgenic lines. (A) *Pf*RNF1-GFP-BirA expression in the transgenic lines. Lysates of mixed asexual blood stage (ABS) of WT NF54 and line *Pf*RNF1-*pfama1*-GFP-BirA and purified immature gametocytes of WT NF54 and line *Pf*RNF1-*pffnpa*-GFP-BirA were immunoblotted with mouse anti-GFP antibodies to detect the *Pf*RNF1-GFP-BirA (∼ 280 kDa). Rabbit antisera against *Pf*39 (39kDa) was used as a loading control. (B) Localization of *Pf*RNF1-GFP-BirA in the transgenic lines. Methanol-fixed schizonts (SZ) of *Pf*RNF1-*pfama1*-GFP-BirA and gametocytes (GC) of stages II-V of *Pf*RNF1-*pffnpa*-GFP-BirA were immunolabeled with mouse anti-GFP antibodies to detect *Pf*RNF1-GFP-BirA (green). SZ was counter labeled with rabbit anti-*Pf*39 antibody and gametocytes with rabbit anti-*Pf*s230 antisera (red); nuclei were highlighted by Hoechst 33342 nuclear stain (blue). Bar, 5 μm. (C) Detection of biotinylated proteins in the transgenic lines. Synchronized ring stages of line *Pf*RNF1-*pfama1*-GFP-BirA and immature gametocyte (imGC) of line *Pf*RNF1-*pffnpa*-GFP-BirA were treated with 50 µM biotin for 24 h. Lysates were immunoblotted with streptavidin coupled to alkaline phosphatase. Lysates of biotin-treated WT NF54 served as negative control. (D) Venn diagram depicting numbers of biotinylated proteins. Ring stages and immature gametocytes were treated with biotin as described above and streptavidin bead-purified biotinylated proteins were purified and subjected to mass spectrometry for identification. Results (A-C) are representative of three independent experiments.

Protein biotinylation in the *Pf*RNF1-*pfama1*-GFP-BirA and *Pf*RNF1-*pffnpa*-GFP-BirA lines was verified by treating ring stages and gametocytes, respectively, with 50 µM biotin for 24 h. Lysates were prepared and subjected to Western blot analysis using streptavidin conjugated to alkaline phosphatase. The blots show multiple bands of potential biotinylated proteins including a band running at approximately 280 kDa, representing biotinylation of *Pf*RNF1-GFP-BirA (Fig. 2C). No prominent bands could be seen in lysates of biotin-treated WT NF54 parasites.

BioID analyses were subsequently employed to analyze the *Pf*RNF1 interactomes in ABSs and gametocytes. For this, rings and immature gametocytes of the *Pf*RNF1-*pfama1*-GFP-BirA and *Pf*RNF1-*pffnpa*-GFP-BirA lines were treated with biotin as described above, and equal amounts of parasites per sample were harvested. Three independent samples were collected from each line. Mass spectrometric analysis was performed on streptavidin-purified protein samples with three technical replicas for each sample. Following exclusion of proteins with signal peptides, which would be expected to follow the secretory pathway, mass spectrometry identified 118 biotinylated proteins in ABSs and 217 proteins in immature gametocytes with 97 (41%) hits shared between the ABSs and immature gametocytes samples (Fig. 2D, Table S1).

### *Pf*RNF1 interacts with proteins involved in RNA-binding and proteasome activity

To gain information on the putative functions of the *Pf*RNF1 interactors, gene ontology (GO) enrichment analyses were performed. Due to the major expression of *Pf*RNF1 in immature gametocytes, we primarily focused on the interaction network in these stages. GO term analysis of 217 proteins identified in immature gametocytes, resulted in biological processes such as translation, macromolecule biosynthesis and genes expression (Fig. 3). Cellular components included particularly the CCR4-NOT core complex with proteins such as CAF1, CAF40, NOT1-G, NOT1, and NOT2 being present. Further, ribosomal proteins, P-bodies, components of the proteasome regulatory particle complex with proteins such as RPN2, RPT1, RPT3, RPT5 and RPT6, and components of the ribonucleoprotein complexes like CITH, PABP1, eEF2, and Puf1 were identified (Fig. 3). In addition, molecular functions of *Pf*RNF1 linked to the proteasome activation, RNA-binding as well as translational initiation were significantly enriched (Fig. 3). To be emphasized are various zinc finger proteins such as PF3D7_1205500, PF3D7_1220000, PF3D7_0927200, PF3D7_0522900 and PF3D7_0602000 (Table S1) which contain at least a CCCH-zinc finger domain postulated to interact with RNA (29).

**Figure 3:**
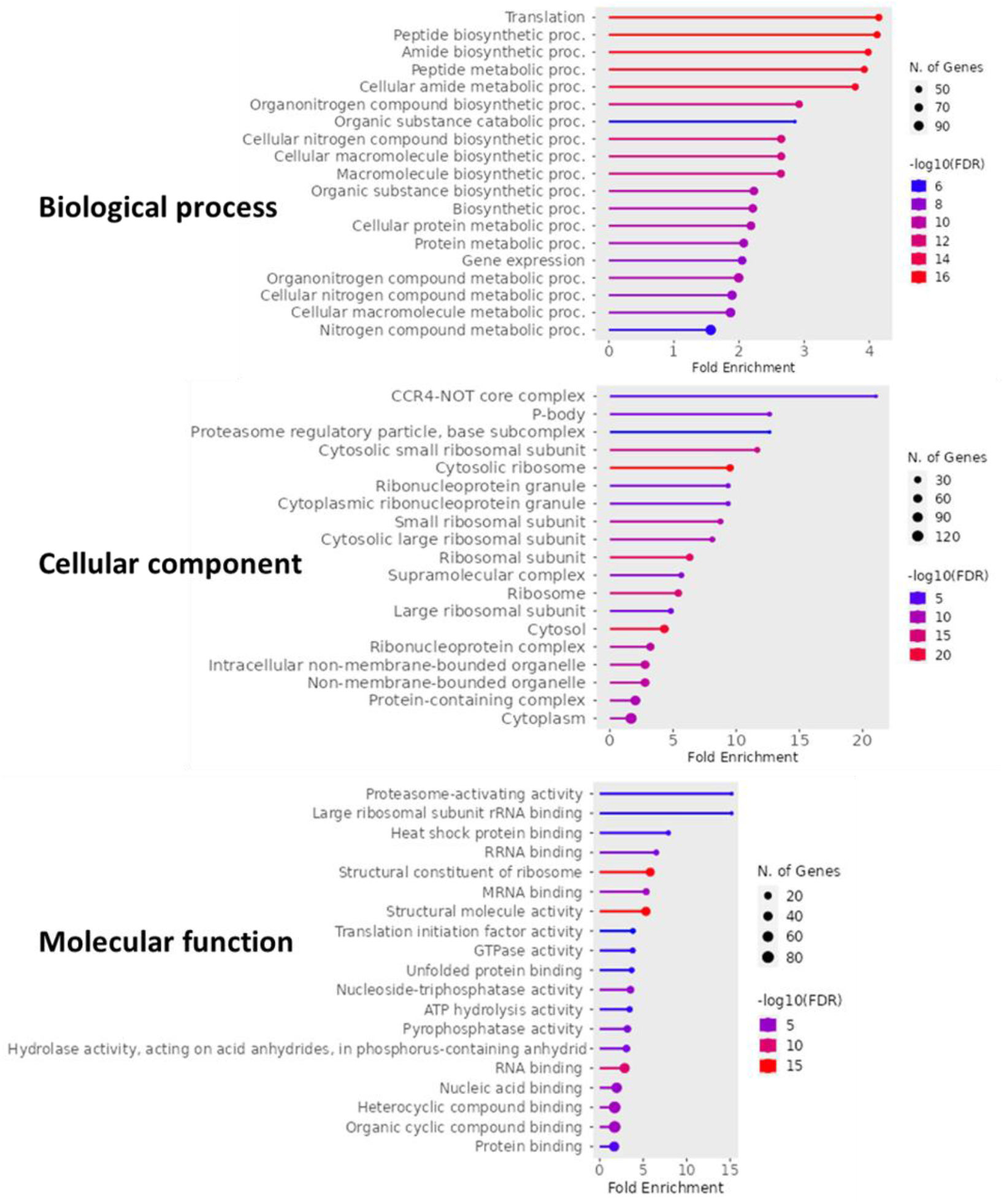
Gene ontology analysis of the *Pf*RNF1 interactome in immature gametocytes. GO enrichment analysis of potential *Pf*RNF1 interactors in immature gametocytes by ShinyGO 0.77 at P < 0.05 reveals the enriched GO terms based on biological process, cellular component and molecular function.

The GO term analyses were also performed on the 118 *Pf*RNF1 interactors in ABS parasites. Biological processes and cellular components identified for the ABS-specific interactors were similar to the ones identified in the gametocyte-specific interactome. Molecular functions of ABS interactors of *Pf*RNF1 included protein-, RNA-, and GTP-binding (Fig. S3).

Furthermore, STRING analyses were performed. The 217 potential *Pf*RNF1 interactors in immature gametocytes formed three major clusters composed of ribosomal proteins, proteasomal proteins and proteins associated with glycolysis were identified (Fig. 4A). KEGG pathway analysis assigned the *Pf*RNF1 interactors to four main cellular activities; ribosomal functions, RNA transport, RNA degradation, and proteasomal activities (Fig. 4B). The STRING pathway analysis of the ABS sample assigned the 118 *Pf*RNF1 interactors mainly to ribosomal proteins (Fig. S4).

**Figure 4:**
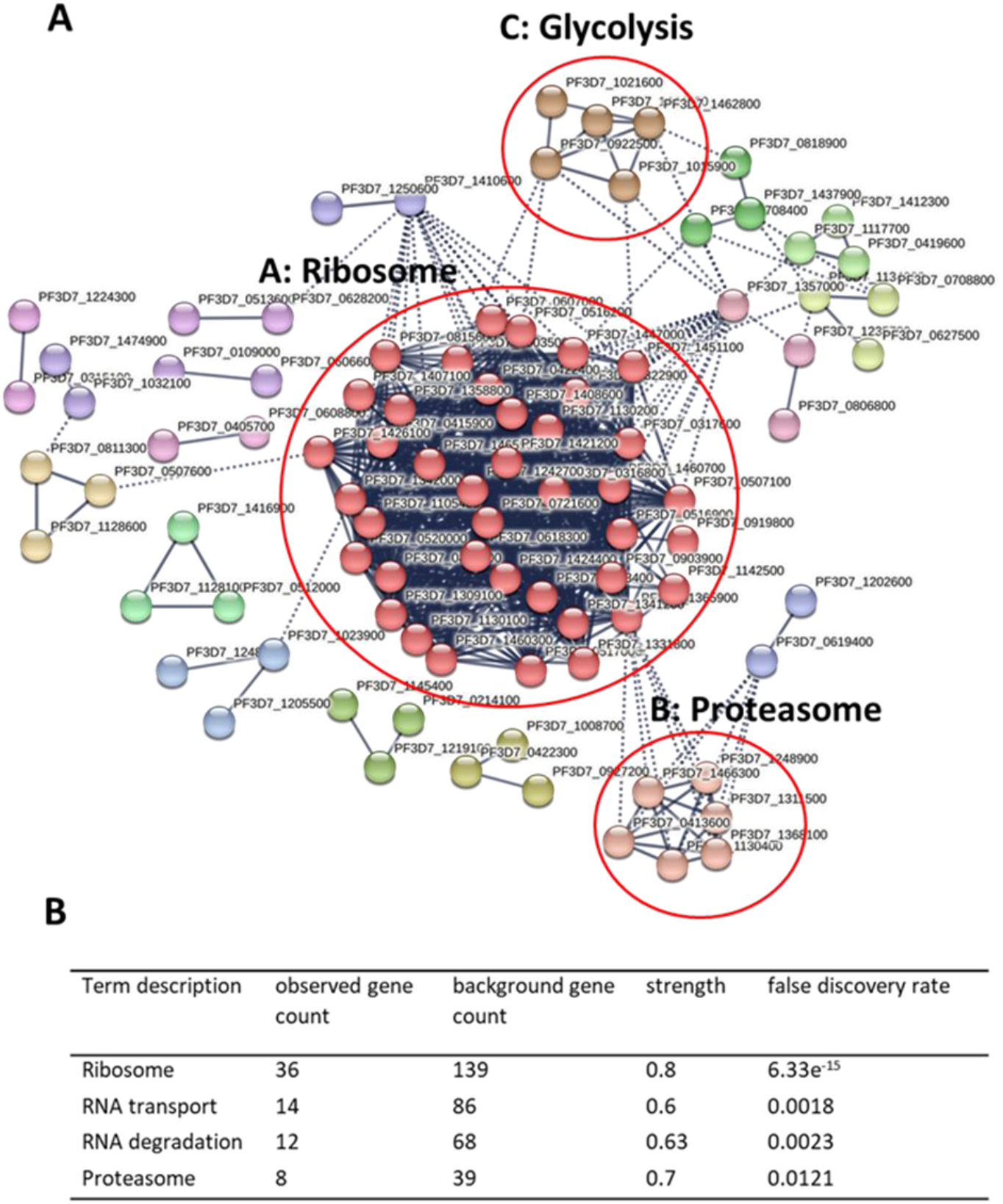
Functional network analysis of the *Pf*RNF1 interactome in immature gametocytes. (A) STRING analysis of the 217 *Pf*RNF1 interactors was assessed using the STRING database (Version 11.5). With highest interaction confidence of 0.9, a network of potential *Pf*RNF1 interactors was generated, where line thickness signifies the strength of data support. Disconnected nodes were not shown in the network. A Markov Clustering (MCL) algorithm was employed to create the possible clusters with an inflation parameter of 3. Three main clusters were identified as proteins of ribosome (Cluster A), proteasome (Cluster B) and the glycolysis pathway (Cluster C). (B) KEGG analysis of the 217 *Pf*RNF1 interactors. Terms describing the KEGG pathway were sorted based on the observed gene count in the whole network.

Interactors of *Pf*RNF1, which were present in both ABSs and gametocytes, also included mostly ribosomal proteins and other RNA-binding proteins associated to ribonucleoprotein complexes like stress granules, the CCR4-NOT core complex as well as proteins associated to the proteasome or heat shock proteins (Table S1).

### *Pf*RNF1 interacts directly with the RNA-binding protein Pumilio-1 (Puf1)

In a last step of our analysis, we investigated the protein-protein interaction of *Pf*RNF1 with its putative interactor Puf1, an RNA-binding protein earlier shown to play an important role in gametocyte differentiation and maintenance (30). For this, a transgenic line expressing HA-tagged Puf1 was generated, using vector pSLI-HA-glmS (Fig. S5A; (28)). Integration of the vector in the *puf1* locus was verified by diagnostic PCR (Fig. S5B) and the expression of HA-tagged Puf1 in ABSs and immature gametocytes of the Puf1-HA-glmS line was demonstrated via Western blot analysis, using anti-HA antibody (Fig. S5C).

To demonstrate the protein-protein interactions between *Pf*RNF1 and Puf1, immature gametocytes of the Puf1-HA-glmS line were purified and the HA-tagged Puf1 was immunoprecipitated, using anti-HA antibody. The precipitated proteins were subjected to Western blotting. Immunoblotting with anti-HA antibody confirmed the precipitation of Puf1 running at the expected molecular weight of 227 kDa, while immunoblotting with anti-*Pf*RNF1 antisera revealed a protein band indicative of *Pf*RNF1 (Fig. 5A). No signal was detected when anti-*Pf*39 antisera was used as control (Fig. 5A). As additional control, we used immature gametocytes of WT NF54 for the immunoprecipitation with anti-HA antibody, which resulted in no detection of either Puf1 or *Pf*RNF1 (Fig. 5B).

**Figure 5:**
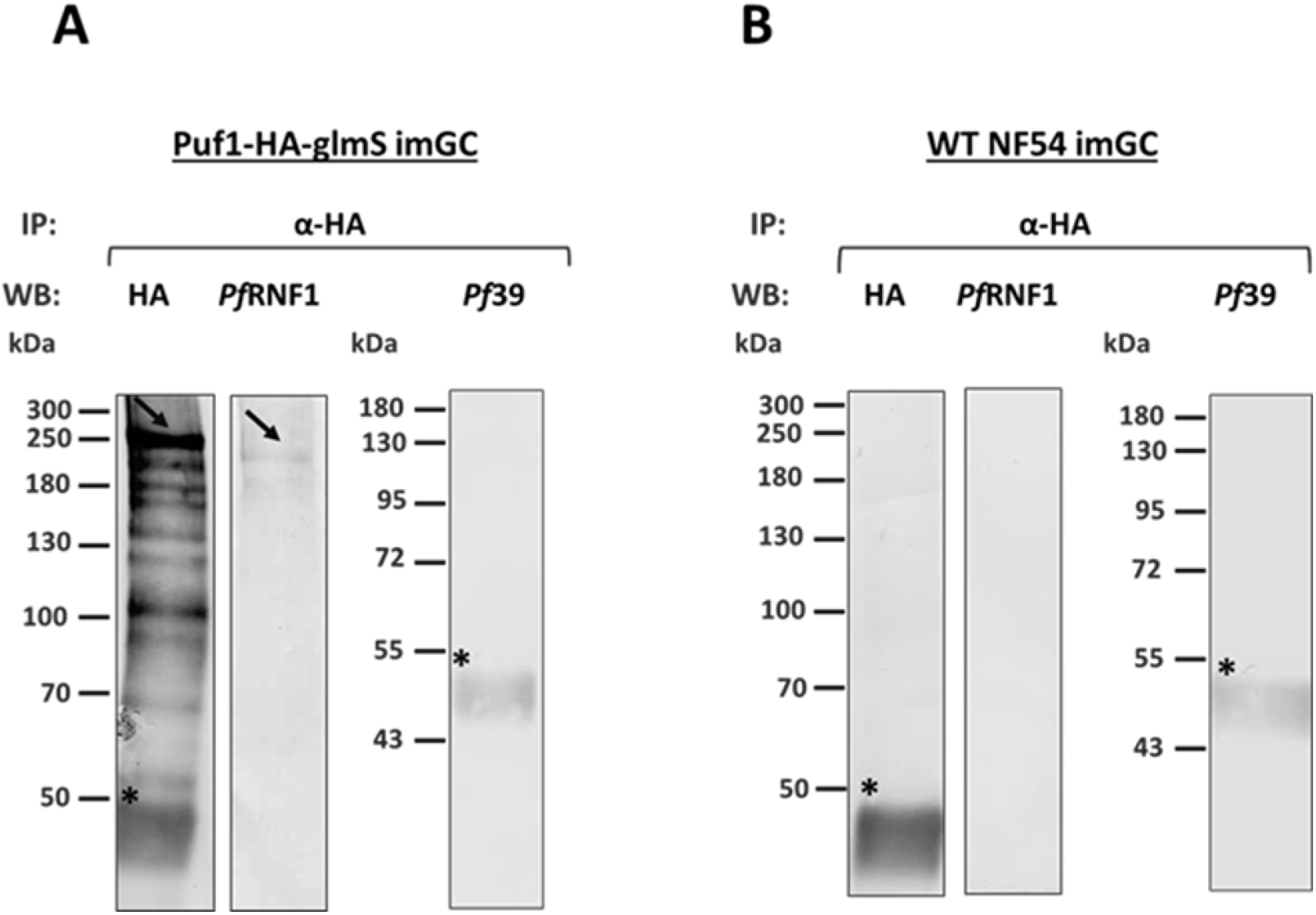
Protein-protein interaction analysis of *Pf*RNF1 with the RNA-binding protein Puf1. (A) Co-immunoprecipitation assays were performed on lysates of immature gametocytes of line Puf1-HA-glmS, using polyclonal rabbit anti-HA antibodies, followed by immunoblotting with mouse anti-*Pf*RNF1 antisera or rabbit anti-HA antibody to detect the precipitated proteins. (Expected sizes: Puf1-HA: ∼227 kDa; *Pf*RNF1: ∼ 180 kDa). Rabbit anti-*Pf*39 antisera was used as a negative control (∼39 kDa). (B) Co-immunoprecipitation on WT NF54 immature gametocytes, following the same procedure, was performed as negative control. Black arrows indicate the precipitated proteins and asterisks show bands corresponding to the precipitation antibody. Results (A, B) are representative of two independent experiments.

## DISCUSSION

Life-cycle progression of *P. falciparum* in the human host and the mosquito vector requires rapid protein turnover activities, and this is mediated by a very tight regulation of transcription, translation and proteolysis. While in recent years, an increasing number of studies unveiled the manifold mechanisms of gene regulation in the blood stages of malaria parasites, not much is known about other processes of proteostasis in these stages. In this study we characterized a putative *P. falciparum* E3 ligase of *P. falciparum* gametocytes named *Pf*RNF1 to unveil its function during gametocyte development.

*Pf*RNF1 belongs to the ring finger proteins (RFPs), a subgroup of zinc finger proteins with diverse functions in transcription, RNA transport, signal transduction, and ubiquitination (31, 32). In contrast to other zinc finger proteins, RFPs mainly execute their function via protein-protein interactions. RFPs are characterized by cysteine-rich domains coordinated by two zinc ions and a RING motif consensus sequence of C-X2-C-X(9-39)-C-X(1-3)-H-X(2-3)-C-X2-C-X(4-48)-C-X2-C (29). Best-studied are RFPs that act as E3 ligases and mediate the transfer of ubiquitin to target proteins thereby initiating their degradation by the UPS (29).

*Pf*RNF1 has originally been described by us in 2017 following a comparative transcriptomics screen in *P. falciparum* gametocytes (24). It possesses a C-terminal RING domain which shows a significant homology with the human E3 ligase Praja-1 known to mediate protein degradation by the UPS (33). *Pf*RNF1 was upregulated in gametocytes following treatment with the histone deacetylase inhibitor Trichostatin A. We therefore postulated that *Pf*RNF1 is a potential HDAC-regulated E3 ligase involved in the UPS during gametocyte development.

We now aimed to gain insights into the role of *Pf*RNF1 by BioID approaches. Initially, we confirmed the expression of *Pf*RNF1 in the blood stage cytoplasm and nucleus with peak expression during gametocyte development. We further demonstrated that *Pf*RNF1 is particularly expressed in male gametocytes, as has been previously indicated by comparative transcript analysis (34). To determine the interaction profile of *Pf*RNF1 and elucidate its physiological role in the parasite, we generated two transgenic lines expressing a *Pf*RNF1-GFP-BirA-fusion protein in either ABSs or gametocytes to be used for BioID analysis, which allowed for the identification of *Pf*RNF1 interactome and proximal proteins. We identified 118 proteins in ABSs and 217 proteins in immature gametocytes as potential interactors of *Pf*RNF1. The high number of hits was identified in immature gametocytes as compared to the ABSs. Ninety-seven hits were shared in both the ABSs and gametocytes. Due to the high abundance of *Pf*RNF1 in immature gametocytes and the fact that most interactors were identified in these stages, we focused our proteomics analyses on the gametocyte biology.

GO enrichment, KEGG, STRING and functional analysis of the identified interactors highlighted the involvement of *Pf*RNF1 with the proteasome complex due to its interaction with components such as RPN2, RPN11, RPT1, RPT3 and RPT6. These results confirm a role of *Pf*RNF1 in the UPS, in accord with its high homology with the human E3-ubiquitin ligase Praja-1 (24) and its previous annotation as an UPS component (20).

Although proteins of the UPS were identified as part of *Pf*RNF1 interactome, the majority of its interactors were associated to translation and included ribosomal proteins and other RNA-binding proteins important for protein synthesis. Ribosomal proteins represented the highest group and several studies have pointed to a major role of the UPS in ribosome quality control whereby the status of nascent polypeptide chain translation is monitored by the ribosome-associated quality control system. When the system detects defects in translation, the nascent peptide chain and the mRNAs associated with it are directed for degradation by the UPS (15). The degradation of the polypeptide takes place following their modification with ubiquitin by E3 ubiquitin ligases as has been shown for ZNF598 and LTN1 in other organisms (35, 36). It is likely that *Pf*RNF1 may play a similar role in *P. falciparum* due to its high association with ribosomal proteins.

RNA-binding proteins such as proteins of the CCR4-NOT core complex like CAF1, CAF40, NOT1-G, NOT1 and NOT2 were also identified. The CCR4-NOT complex is a conserved large multifunctional assembly of proteins which plays an important role in mRNA decay (37). Some members of this complex e.g., CNOT4 acts as an RNA-binding E3 ligases which exhibit both E3 ligase and RNA-binding activity (38), similar to the predicted functions of *Pf*RNF1. Noteworthy, a high variety of proteins involved in translational control were previously identified in malaria parasites (2) and in *P. yoelii, Py*NOT1-G and *Py*CCR4-1 are members of the CCR4-NOT complex which have been shown to play an important role in gametocyte development and transmission by regulating mRNAs important for the processes (8).

Another group of proteins identified in the *Pf*RNF1 interactome are proteins associated with translational repression such as CITH, PUF1, DOZI and PABP1. These proteins associate with non-translated mRNAs to form a mRNP complex in the form of stress granules or P-bodies. The repressed mRNAs are stored in these stress granules and released at a later time point to be either introduced to the protein synthesis machinery or to the proteasome for degradation. In *P. berghei* and *P. falciparum*, it was demonstrated that transcripts of the parasite important for mosquito midgut stage formation are synthesized and stored in female gametocytes granules where they are translationally repressed by binding to regulatory RNA-binding proteins like CITH and DOZI. The repression is only released after gametocyte activation to promote zygote to ookinete formation (10, 12, 13). Many UPS-associated proteins have been found in these mRNP complex granules for example the ubiquitin ligase Roquin and MEX3 are components of granules (39–41). It is possible that *Pf*RNF1 also associates with these granules.

Another interesting finding in this study was the identification of some zinc finger proteins as *Pf*RNF1 interactors in both ABSs and immature gametocytes, all of which have been reported to contain at least one CCCH zinc finger domain which can interact with RNA (29). Among these CCCH-zinc finger interactors, *Pf*ZNF4 was identified. *Pf*ZNF4 was recently shown by our group to regulate male enriched transcripts in gametocytes (42). In accord with these findings, *Pf*RNF1 is particularly found in male gametocytes.

It was surprising to identify a strong association of *Pf*RNF1 with RNA-binding proteins by BioID analyses, since no defined RNA-binding domain was identified in the protein. However, other proteins such as ARIH2 also possess E3 ligase and RNA-binding properties but lacks a well-defined RNA binding domain (15), suggesting that may also exert dual activities as E3 ligase and RNA binder.

In a recent study, *Pf*RNF1 was identified in a protein interaction network with md1 which is an important determiner of male gametocyte fate in *P. falciparum* (43). It was also interesting to see that the authors identified proteins associated to the CCR4-NOT complex as well as proteins associated with translational repression in the interaction network which therefore confirms our data on the involvement of *Pf*RNF1 in these processes.

Our combine data therefore indicate that *Pf*RNF1 is a multifunctional RBUL protein which links the UPS with RNA-binding proteins to regulate the post-transcriptional machinery in the malaria parasite *P. falciparum*.

## MATERIALS AND METHODS

### Antibodies

The following antibodies were used in the study: rat anti-HA (Roche, Basel, Switzerland), rabbit anti-HA (Sigma-Aldrich), mouse anti-GFP (Roche, Basel, Switzerland), rabbit anti-*Pf*s230 (BioGenes, Berlin, Germany), mouse anti-*Pf*39 (24), rabbit anti-*Pf*39 (Davids Biotechnology, Regensburg, Germany), rabbit *Pf*s25 (ATCC, Manassas, USA), rabbit anti-*Pf*MSP1 (ATCC, Manassas, USA). The mouse anti-sera against *Pf*RNF1 fragment 1 (RP1) was produced previously (24) while mouse anti-sera against *Pf*RNF1 fragment 2 (RP2) was produced in this study. The following dilutions were used for IFA: rabbit anti-*Pf*s230 (1:500), mouse anti-*Pf*RNF1-RP1 (1:20), mouse anti-*Pf*RNF1-RP2 (1:20), mouse anti-GFP (1:200). For Western blot analysis the following dilutions were used: rat anti-HA (1:500), mouse anti-*Pf*39 (1:1000), rabbit anti-*Pf*39 (1:10000), mouse anti-GFP (1:1000), and anti -*Pf*RNF1-RP2 (1:500).

### Parasite culture

The high gametocyte producing strain *P. falciparum* NF54 (termed WT NF54) was used in all experiments as well as for the generation of transgenic lines. The parasites were cultured *in vitro* in RPMI 1640/HEPES medium (Gibco, Thermo Scientific Waltham, USA) containing 10% heat-inactivated human serum and A^+^ erythrocytes at 5% hematocrit (44). 50 μg/ml hypoxanthine (Sigma Aldrich, Taufkirchen, Germany) and 10 μg/ml gentamicin (Gibco,

Thermo Scientific Waltham, USA) were also added to the cell culture medium as supplements. Cultivation was performed with a gas mixture of 5% O_2,_ 5% CO_2,_ 90% N_2_ at a temperature of 37°C. To obtain synchronized cultures, they were treated with 5% sorbitol as described (45). Human erythrocyte concentrate and serum were purchased from the transfusion medicine department of the University Hospital Aachen, Germany. The work on human blood was approved by the University Hospital Aachen Ethics commission (EK007/13) and serum samples were pooled and the donors remained anonymous.

### Generation of mouse PfRNF1-RP2 antisera

Recombinant peptides, corresponding to a portion of *Pf*RNF1 (RP2; Fig. 1A) was expressed as a maltose-binding fusion protein using the pMAL™c5X-vector (New England Biolabs, Ipswich, USA). To this end, the coding sequence was amplified using gene-specific primers (for primer sequences, see Table S2) and the recombinant protein was expressed in *E. coli* BL21 (DE3) RIL cells following the manufacturer’s protocol (Invitrogen, Karlsruhe, Germany). The protein was then purified by affinity-purification using an amylose beads according to the manufacturer’s protocol (New England Biolabs, Ipswich, USA) and the concentration determined by Bradford assay. 100 µg of pure protein emulsified in Freund’s incomplete adjuvant (Sigma Aldrich, Taufkirchen, Germany) was injected subcutaneously to six weeks old female NMRI mice (Charles River Laboratories, Wilmington, USA). This was followed by a boost after 4 weeks and 10 days after the boost, mice were anesthetized through intraperitoneal injection of a mixture of ketamine and xylazine according to the manufacturer’s protocol (Sigma Aldrich, Taufkirchen, Germany), and immune sera were collected via heart puncture. Immune sera from three immunized mice were pooled; sera of three non-immunized mice (NMS) were used as negative control. Experiments in mice were approved by the animal welfare committee of the District Council of Cologne, Germany (ref. no. 84-02.05.30.12.097 TVA).

### Generation of transgenic parasite lines Generation of the *Pf*RNF1-GFP-BirA parasite lines

To investigate the *Pf*RNF1 interaction network, the BioID method was used (46) in which the *Pf*RNF1-BirA lines were generated that episomally overexpress the full *Pf*RNF1 fused to a biotin ligase (Bir A). To achieve this, full length sequence of *Pf*RNF1 was cloned into the vector pARL-GFP-BirA (for primer sequences, see Table S2), under the control of the *ama1* (Apical membrane antigen 1) promoter, which is mainly expressed in the ABSs and the gametocyte-specific *fnpa* promoter, that is mainly expressed in the sexual blood stages using the pARL-*pfama1*-GFP-BirA and pARL-*pffnpa*-GFP-BirA vectors respectively as described before (28). The constructed plasmids were then used to transfect parasites as described above. After 21 days parasites were visible and the selection of parasites episomally overexpressing the proteins was done by treatment with medium containing 4nM WR99210 (Jacobus Pharmaceutical Company, USA).The successful uptake of the vector was confirmed by diagnostic PCR (Fig. S2B; for primer sequences, see Table S2) The house keeping gene aldolase was used as control and was amplified using *pffbpa* specific primers as described before (28).

### Generation of Puf1-HA-glmS parasite line

Parasite line in which Puf1-was tagged with a hemagglutinin (HA) tag and a glmS-ribozyme was generated (47) by using the pSLI-HA-glmS vector (Fig. S5A; kindly provided by Dr. Ron Dzokowski, the Hebrew University of Jerusalem). To achieve this, we modified the plasmid to contain a homology block from the 3’ end of the Puf1 gene without the stop codon (for primer sequences, see Table S2) thereby enabling the coding region for Puf1 to be fused at the 3’-region to a HA-encoding sequence followed by the glmS-ribozyme sequence in the vector. Parasites were transfected with the modified plasmid and WR99210 was added to a final concentration of 4 nM, starting at 6 h after transfection to select for integrated parasites. WR99210-resistant parasites appeared at ∼21 days post-transfection and they were treated with medium supplemented with 400 μg/ml G418 and vector integration was verified by diagnostic PCR (for primer sequences, see Table S2). Vector integration for the parasite line was successful and the tagged protein could be detected using HA-antibodies (Fig. S5B, C).

### Co-immunoprecipitation Assay

Percoll purified immature gametocytes of WT NF54 and the Puf1-HA-glmS line were lysed in RIPA buffer (150mM NaCl, 1% Triton X-100, 0.5% sodium deoxycholate, 0.1% sodium dodecyl sulphate, 50mM Tris in distilled water). The lysates were incubated on ice for 15 min. Three sessions of sonication were applied to each sample (30sec/ 50% and 0.5 cycles). After centrifugation (16,000 x g for 10 min at 4°C), the supernatant was incubated with 5% v/v pre-immune rabbit sera and 20 μl of protein G-beads (Roche) for 1 h on a rotator at 4°C. After centrifugation (3500 x g for 5 min at 4°C), the supernatant was incubated for 1 h at 4°C with 5% v/v polyclonal rabbit antisera against HA. Afterwards, a volume of 30 μl protein G-beads was added and kept on rotation overnight at 4°C. Following centrifugation (3500 x g for 5 min at 4°C), beads were first washed with ice cold RIPA buffer and then with PBS for five times. The beads were finally resuspended in an equal volume of loading buffer. The samples were then subjected to Western blotting as described below.

### Western Blotting

ABS parasites of the WT NF54, the *Pf*RNF1-HA-glmS line, the *Pf*RNF1-*pfama1*-GFP-BirA, and the *Pf*RNF1-*pffnpa*-GFP-BirA lines were harvested from mixed or synchronized cultures, while gametocytes were enriched by Percoll purification. Parasites were released from iRBCs with 0.015% w/v saponin/PBS for 10 min at 4°C, then washed with PBS, and resuspended in lysis buffer (0.5%v/v Triton X-100, 4% w/v SDS, 50% v/v 1xPBS) which was supplemented with protease inhibitor cocktail; 5x SDS-PAGE loading buffer containing 25 mM DTT was added to the lysates, samples were heat-denatured for 10 min at 95°C and separated using the SDS-PAGE. Parasite proteins separated by gel electrophoresis were transferred to Hybond ECL nitrocellulose membrane (Amersham Biosciences, Buckinhamshire, UK) as per the manufacturer’s protocol. The membranes were incubated in Tris-buffered saline containing 5% w/v skim milk, pH 7.5 to prevent Non-specific binding, then followed by immune recognition overnight at 4°C using polyclonal mouse anti-*Pf*39 antisera, mouse anti-GFP antibody, mouse anti-*Pf*RNF1 or rat anti-HA antibody. After washing, membranes were incubated with the respective alkaline phosphatase-conjugated goat secondary antibody (Sigma-Aldrich) for 1 h at room temperature (RT). Biotinylated proteins were directly labeled using alkaline phosphatase-coupled streptavidin (Sigma-Aldrich). The blots were developed in a solution of nitroblue tetrazolium chloride (NBT) and 5-brom-4-chlor-3-indoxylphosphate (BCIP; Merck, Darmstadt, Germany) for 5–30 min at RT. Blots were scanned and processed using the Adobe Photoshop CS software. Band intensities were measured using the ImageJ program version 1.51f.

### Indirect Immunofluorescence Assay

Cultures containing mixed ABSs and gametocytes of the *Pf*RNF1-HA-KD, the *Pf*RNF1-*pfama1*-GFP-BirA, and the *Pf*RNF1-*pffnpa*-GFP-BirA parasite lines were air-dried as cell monolayers on glass slides and then fixed in a methanol bath at -80°C for 10 min. The fixed cells were sequentially incubated in 0.01% w/v saponin/0.5% w/v BSA/PBS and 1% v/v neutral goat serum (Sigma-Aldrich)/PBS for 30 min at RT to facilitate membrane permeabilization and blocking of non-specific binding. Thereafter, the preparations were incubated with mouse anti-*Pf*RNF1, rat anti-HA or mouse anti-GFP antibody, diluted in 0.5% w/v BSA/PBS for 2 h at 37°C. Washing steps were carried out, followed by binding of the primary antibody and a detection step facilitated by the incubation with Alexa Fluor 488-conjugated goat anti-rat or anti-mouse secondary antibody (Thermo Fisher Scientific, Waltham, MA, USA). Alexa Fluor 594-conjugated streptavidin was used (Thermo Fisher Scientific, Waltham, MA, USA) to immunolabel biotinylated proteins. Mouse antisera directed against *Pf*39 or *Pf*92 were used to highlight the ABSs, while rabbit antisera directed against *Pfs*230 were used to highlight gametocytes, followed by incubation with polyclonal Alexa Fluor 488-or 594-conjugated goat anti-mouse or anti-rabbit secondary antibodies (Invitrogen Molecular Probes; Eugene, OR, USA). Alternatively, the ABSs were stained with 0.01% w/v Evans Blue (Sigma-Aldrich; Taufkirchen, Germany)/PBS for 3 min at RT followed by 5 min washing with PBS. The parasite nuclei were highlighted by incubation with Hoechst 33342 nuclear stain (Invitrogen) for 5 min at RT. After washing with PBS, cells were mounted with anti-fading solution AF2 (CitiFluorTM, Hatfield, PA, USA), and sealed with nail polish. Specimen were examined with a Leica DM 5500 B microscope, and digital images were processed using the Adobe Photoshop CS software.

### Preparation of samples for BioID analysis

Highly synchronized ring stage *P. falciparum* cultures of the *Pf*RNF1-*pfama1*-BirA parasite line and Percoll-enriched immature gametocyte cultures of the *Pf*RNF1-*pffnpa*-BirA parasite lines were treated for 24 h with biotin at a final concentration of 50 µM to induce proximal biotinylation of proteins by the overexpressing *Pf*RNF1 tagged BirA ligase. Following treatment, the erythrocytes were lysed with 0.05% w/v saponin/PBS and the released parasites were resuspended in 100-200 µl binding buffer (Tris-buffered saline containing 1% v/v Triton X-100 and protease inhibitor) in a low protein binding Eppendorf tube and the samples were then sonicated on ice (2x 60 pulses at 30% duty cycle). Another 100 µl cold Tris-buffered saline was added and sonication was again performed. After centrifugation (5 min, 16,000xg, 4°C) the supernatant was removed and transferred into another reaction tube and mixed with 100 µl pre-equilibrated Cytiva Streptavidin Mag Sepharose™ Magnet-Beads (Thermo Fisher Scientific, Walham, USA). The samples were then incubated with slow end-over-end mixing at 4°C overnight. Following six washing steps to remove unbound proteins (3x with RIPA buffer containing 0.03% w/v SDS and three times with 25 mM Tris buffer, pH 7.5), biotinylated proteins bound to the beads were eluted by the addition of 40 µl of 1% (w/v) SDS/5 mM biotin in Tris buffer (pH 7.5) and incubation at 95°C for 5-10 min.

### Proteolytic digestion

Processing of samples was done by single-pot solid-phase-enhanced sample preparation (SP3) as described (48, 49). Briefly, the eluted biotinylated proteins were reduced and alkylated, using DTT and iodoacetamide (IAA), respectively. Afterwards, 2 µl of carboxylate-modified paramagnetic beads (Sera-Mag Speed Beads, GE Healthcare), 0.5 μg solids/μl in water was added. After adding acetonitrile to a final concentration of 70% (v/v), samples were left to settle at RT for 20 min. Subsequently, beads were washed twice with 70 % (v/v) ethanol in water and one time with acetonitrile. Beads were resuspended in 50 mM NH_4_HCO_3_ supplemented with trypsin (Mass Spectrometry Grade, Promega) at an enzyme-to-protein ratio of 1:25 (w/w) and the samples were incubated overnight at 37°C. After overnight digestion, acetonitrile was added to the samples at a final concentration of 95% (v/v) and incubated at RT for 20 min. To improve the yield, supernatants derived from this initial peptide-binding step were additionally subjected to the SP3 peptide purification procedure (49) Each sample was then washed with acetonitrile. To recover bound peptides, paramagnetic beads from the original sample and corresponding supernatants were pooled in 2 % (v/v) dimethyl sulfoxide (DMSO) in water and sonicated for 1 min. After 2 min of centrifugation at 12,500 rpm and 4 °C, supernatants containing tryptic peptides were transferred into a glass vial for MS analysis and acidified with 0.1 % (v/v) formic acid.

### Liquid chromatography-mass spectrometry (LC-MS) analysis

LC-MS was performed as described before (28). To this end, tryptic peptides were separated using an Ultimate 3000 RSLCnano LC system (Thermo Fisher Scientific) equipped with a PEPMAP100 C18 5 µm 0.3 × 5 mm trap (Thermo Fisher Scientific) and an HSS-T3 C18 1.8 μm, 75 μm x 250 mm analytical reversed-phase column (Waters Corporation). Mobile phase A was water containing 0.1 % (v/v) formic acid and 3 % (v/v) DMSO. Peptides were separated running a gradient of 2–35% mobile phase B (0.1% (v/v) formic acid, 3 % (v/v) DMSO in ACN) over 40 min at a flow rate of 300 nl/min. The total time of analysis was 60 min including wash and column re-equilibration steps. Column temperature was set to 55°C. Mass spectrometric analysis of eluting peptides was conducted on an Orbitrap Exploris 480 (Thermo Fisher Scientific) instrument platform with a spray voltage of 1.8 kV, funnel RF level of 40, and heated capillary temperature of 275°C. Data were acquired in data-dependent acquisition (DDA) mode targeting the 10 most abundant peptides for fragmentation (Top10). Full MS resolution was set to 120,000 at m/z 200 and full MS automated gain control (AGC) target to 300% with a maximum injection time of 50 ms. Mass range was set to m/z 350–1,500. For MS2 scans, collection of isolated peptide precursors was limited by an ion target of 1 × 105 (AGC target value of 100%) and maximum injection times of 25 ms. Fragment ion spectra were acquired at a resolution of 15,000 at m/z 200. Intensity threshold was kept at 1E4. Isolation window width of the quadrupole was set to 1.6 m/z and normalized collision energy was fixed at 30%. All data were acquired in profile mode using positive polarity. Samples were analyzed in three technical replicates.

### Data analysis and label-free quantification

The DDA raw was acquired with the Exploris 480 and processed using MaxQuant (version 2.0.1) (50, 51) with standard settings and label-free quantification (LFQ) enabled for each parameter group, i.e. the control and affinity-purified samples (LFQ min ratio count 2, stabilize large LFQ ratios disabled, match-between-runs). The data were then quarreled against the forward and reverse sequences of the *P. falciparum* proteome (UniProtKB/TrEMBL, 5,445 entries, UP000001450, release April 2020) and a list of common contaminants. For the identification of peptide, trypsin was set as protease allowing two missed cleavages. Carbamidomethylation was set as fixed and oxidation of methionine as well as acetylation of protein N-termini as variable modifications. Only peptides with a minimum length of 7 amino acids were considered. The peptide and protein false discovery rates (FDR) were set to 1 %. In addition, proteins had to be identified by at least two peptides. Statistical analysis of generated data was performed using Student’s t-test and corrected by the Benjamini–Hochberg (BH) method for multiple hypothesis testing (FDR of 0.01). In addition, proteins in the affinity-enriched samples had to be identified in all three biological replicates and to show at least a two-fold enrichment as compared to the controls.

The datasets of protein hits were again edited by verification of the gene IDs and gene names via the PlasmoDB database (www.plasmodb.org; (52)). Since *Pf*RNF1 has earlier be shown to be expressed in the cytoplasm and nucleus, hits containing signal peptides were removed following their prediction using SignalP-6.0 (https://dtu.biolib.com/SignalP-6). Gene ontology (GO) enrichment and KEGG analysis was performed using ShinyGO 0.77 at P < 0.05 (http://bioinformatics.sdstate.edu/go/, accessed, January 10, 2023). Network analysis was conducted using the STRING database (https://string-db.org/, accessed, January 10, 2023 using default settings and confidence of 0.009.

## Data availability

The mass spectrometry proteomics data have been deposited to the ProteomeXchange Consortium (http://proteomecentral.proteomexchange.org) via the jPOST partner repository (53) with the dataset identifiers PXD040384 for ProteomeXchange and JPST002050 for jPOST.

## Acknowledgments

We thank Olga Papst for the expression of the *Pf*RNF1 recombinant proteins and performing some of the immunofluorescence assays. The authors acknowledge funding by the Deutsche Forschungsgemeinschaft (projects grants NG170/1-1 to CJN and PR905/20-1 to GP and grants PR905/19-1 to GP and TE599/9-1 to ST of the DFG priority programme SPP 2225). JPM and AF received a fellowship from the German Academic Exchange Service (DAAD).

